## Supplemental figure1-5 for "Whole genome sequencing revealed genetic diversity and selection of Guangxi indigenous chickens"

### Supplementary

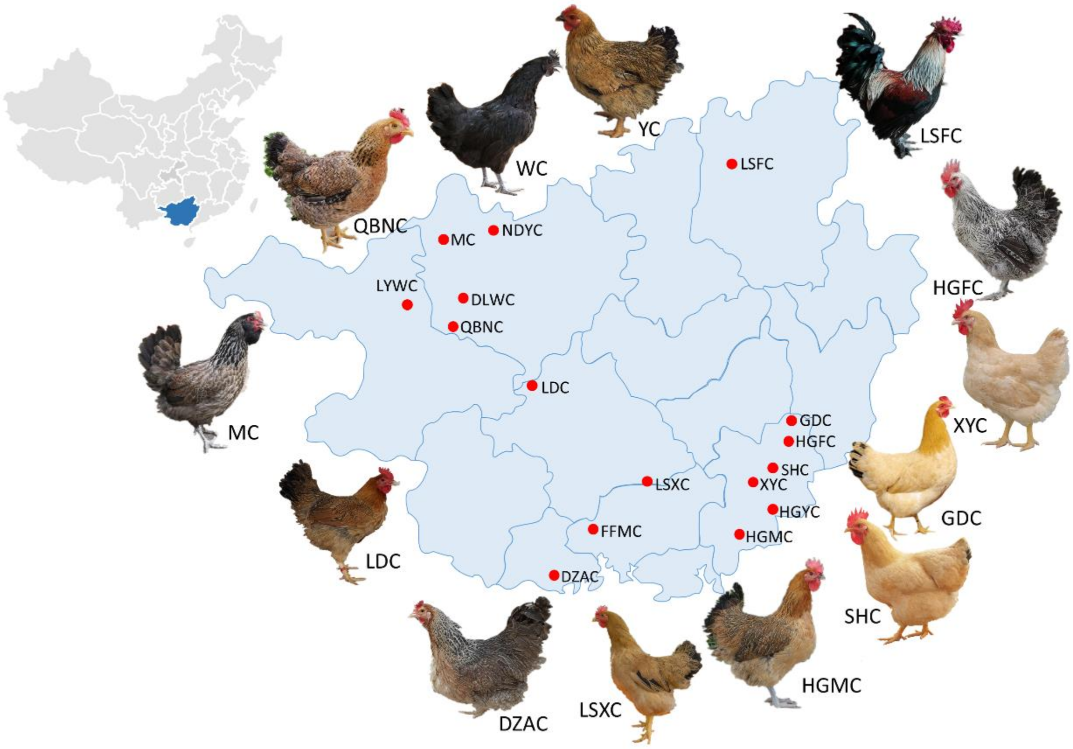


**S1 Fig. Geographic distribution and appearances of typical female chickens.**

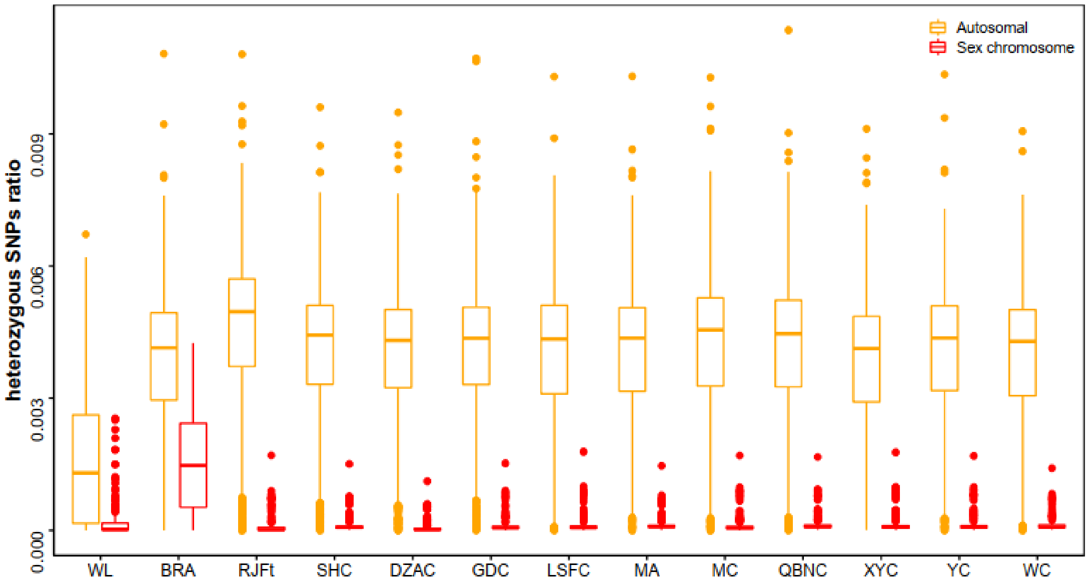


**S2 Fig. Boxplot showing heterozygous SNP rate of autosomes (left) and Z chromosome (right) between each chicken population.**

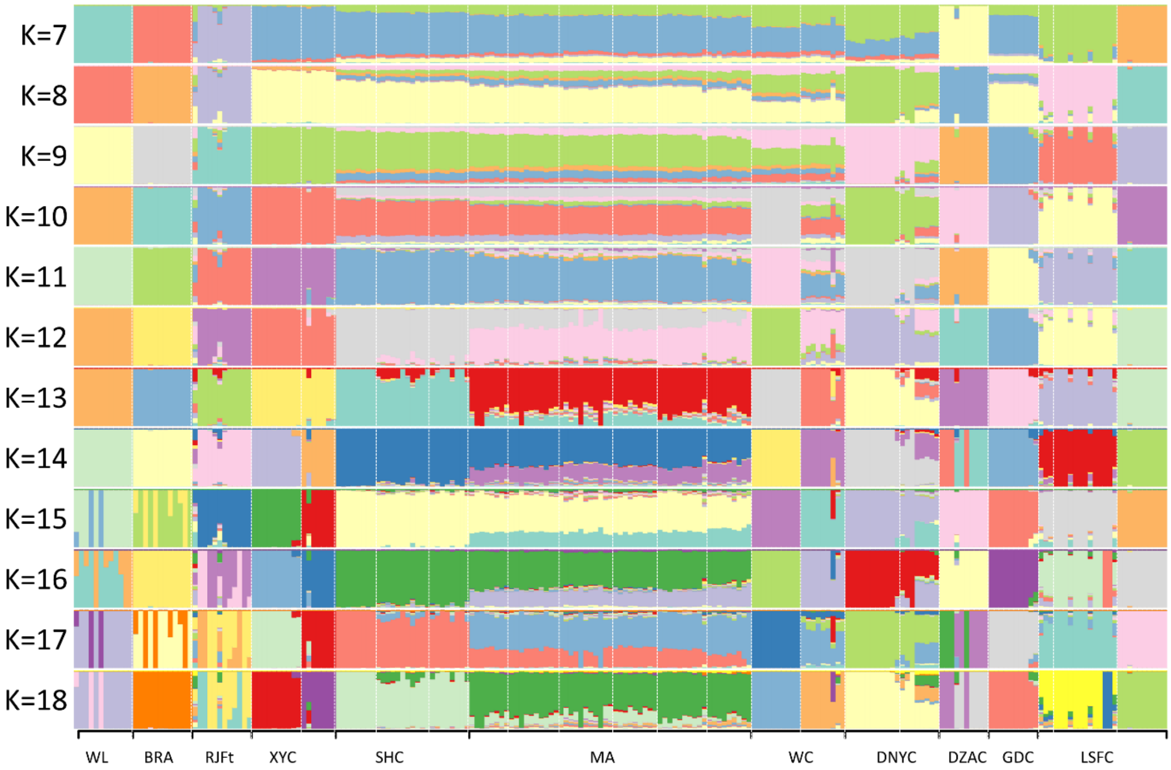


**S3 Fig. Admixture analysis with K values running from 7 to 18.** Each population separated by white dotted line.

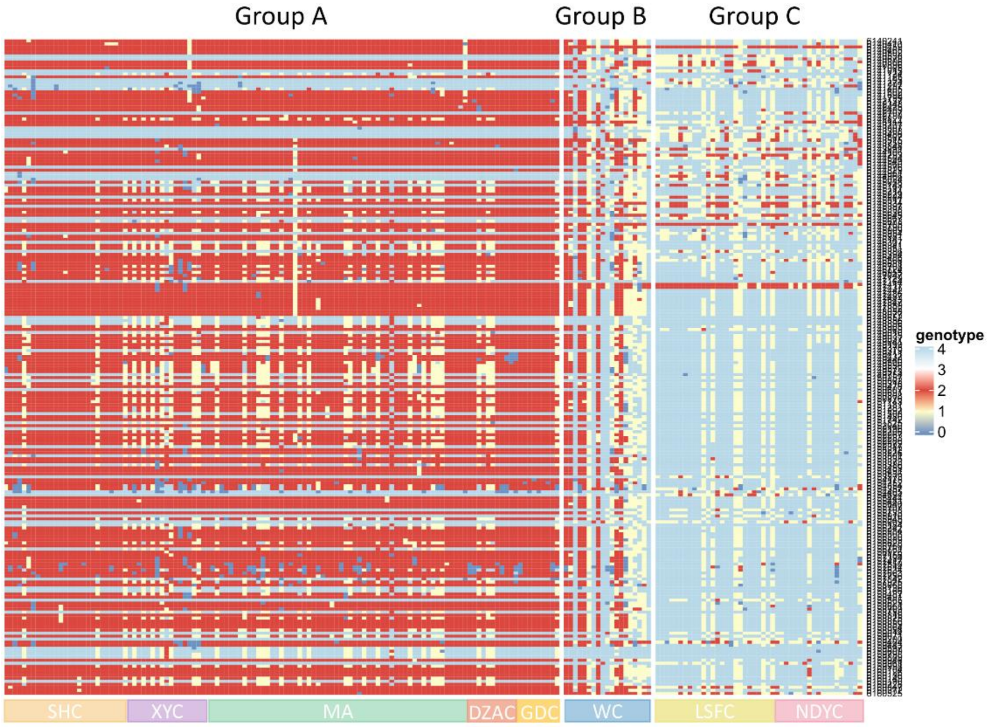


**S4 Fig. The genotype of fixed SNPs in chr24: 6.14Mb~6.18Mb of individuals.** The row represents the SNP position and the column represents the individual. Light blue denotes reference alleles while red indicates alternative homozygous alleles, yellow means heterozygous and dark blue means missing.

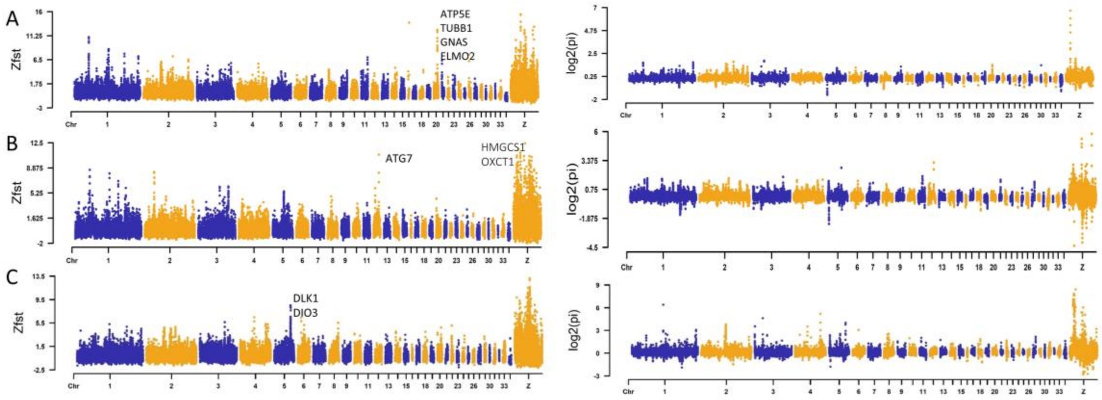


**S5 Fig. ZFst values and Log 2 (pi).** (A) WC and other indigenous population. (B) XYC and SHC. (C) GDC and SHC
